# Phosphorylation of Ribosomal Protein S6 differentially affects mRNA translation based on ORF length

**DOI:** 10.1101/2021.03.18.436059

**Authors:** Jonathan Bohlen, Aurelio A. Teleman

## Abstract

Phosphorylation of Ribosomal Protein S6 (RPS6) was the first post-translational modification of the ribosome to be identified and is a commonly-used readout for mTORC1 activity. Although the cellular and organismal functions of RPS6 phosphorylation are known, its molecular consequences on translation are less well understood. Here we use selective ribosome footprinting to analyze the location of ribosomes containing phosphorylated RPS6 on endogenous mRNAs in cells. We find that RPS6 becomes progressively dephosphorylated on ribosomes as they translate an mRNA. As a consequence, average RPS6 phosphorylation is higher on mRNAs with short coding sequences (CDSs) compared to mRNAs with long CDSs. In particular, ribosomes translating on the endoplasmic reticulum are more rapidly dephosphorylated than cytosolic ribosomes. Loss of RPS6 phosphorylation causes a correspondingly larger drop in translation efficiency of mRNAs with short CDSs than long CDSs. Interestingly, mRNAs with 5’ TOP motifs are translated well also in the absence of RPS6 phosphorylation despite short CDS lengths, suggesting they are translated via a different mode. In sum this provides a dynamic view of RPS6 phosphorylation on ribosomes as they translate mRNAs in different subcellular localizations and the functional consequence on translation.

## INTRODUCTION

A large number of post-transcriptional and post-translational modifications of ribosomal RNA and ribosomal proteins have been discovered (Henras, Soudet et al., 2008, Simsek & Barna, 2017), however the impact of these modifications on ribosomal function is often not understood. While modifications of rRNA are implicated in the biosynthesis, catalytic activity and structural integrity of the ribosome (Granneman & Baserga, 2004, Penzo & Montanaro, 2018), modifications of ribosomal proteins, which include phosphorylation (Gressner & Wool, 1974, Mazumder, Sampath et al., 2003), ubiquitinylation (Higgins, Gendron et al., 2015, Silva, Finley et al., 2015), NEDDylation (Xirodimas, Sundqvist et al., 2008), methylation (Couttas, Raftery et al., 2012), UFMylation (Walczak, Leto et al., 2019) and acetylation (Kamita, Kimura et al., 2011), are functionally less well understood. Nonetheless, these ribosomal protein modifications are of interest because they can be inducible (Gressner & Wool, 1974, Higgins et al., 2015, Mazumder et al., 2003), suggesting they could have regulatory functions.

Inducible phosphorylation of ribosomal protein S6 (RPS6) was discovered in mouse livers after injury (Gressner & Wool, 1974), representing the first known posttranslational modification of the ribosome (Biever, Valjent et al., 2015). RPS6 phosphorylation is placed by S6K upon mTORC1 activation, and by RSK upon ERK activation, making it highly responsive to nutrient availability and growth factor signaling (Meyuhas, 2015). As such, p-RPS6 is widely used as a readout of mTOR pathway activation (Biever et al., 2015, Meyuhas, 2015, Simsek & Barna, 2017). The functional role of p-RPS6 has been elucidated using knock-in mice harboring non-phosphorylatable Rps6 (Ruvinsky, Sharon et al., 2005). Interestingly, these mice show no major growth phenotypes, but instead have impaired glucose homeostasis (Ruvinsky et al., 2005), reduced muscle strength (Ruvinsky, Katz et al., 2009) and impaired compensatory renal hyperthrophy (Xu, Chen et al., 2015). On a cellular level, loss of RPS6 phosphorylation leads to a higher protein synthesis rate (Ruvinsky et al., 2005), smaller cell size (Granot, Swisa et al., 2009, Ruvinsky et al., 2009) and faster proliferation (Ruvinsky et al., 2005). Although the phenotypic consequences of RPS6 phosphorylation at the organismal and cellular levels are clear, the functional consequences of RPS6 phosphorylation at the molecular level are not known. This has been a longstanding open question in the field (Biever et al., 2015, Meyuhas, 2015, Simsek & Barna, 2017). Because of the prominent position of RPS6 phosphorylation on the small ribosomal subunit, one common hypothesis is that it affects some aspect of ribosome activity such as translation initiation or elongation.

Selective ribosome footprinting is a method that identifies the location of specific ribosome sub-populations on endogenous mRNAs in human cells (Bohlen, Fenzl et al., 2020, Schibich, Gloge et al., 2016, Wagner, Herrmannova et al., 2020). This method employs immunoprecipitation to isolate ribosomes bound to specific factors such as initiation factors or chaperones, followed by sequencing of their footprints to identify their locations. Here we adapt this method to look at post-translational modifications of the ribosome. By immunoprecipitating ribosomes phosphorylated on RPS6, we study the location and frequency of phospho-RPS6 ribosomes on endogenous mRNAs in human and mouse cells. We find that ribosomes become progressively dephosphorylated on RPS6 as they move along an mRNA to translate the coding sequence (CDS). This dephosphorylation is rather slow compared to ribosome movement, so that it only becomes appreciable on mRNAs with long coding sequences. We find that RPS6 phosphorylation promotes translation of mRNAs, so that loss of RPS6 phosphorylation mainly causes mRNAs with short CDSs to become translationally repressed. Interestingly, mRNAs containing a 5’ TOP motif defy this general pattern despite being short, and do not require RPS6 phosphorylation for their efficient translation, leading to a relative increase in ribosomal proteins in RPS6 phospho-deficient cells. This work thereby provides a dynamic view of phosphorylation on RPS6 as ribosomes move along mRNAs, as well as a molecular functional outcome for this post-translational modification.

## RESULTS

### Ribosomes with phospho-S6 are depleted from mRNAs with long ORFs

We used selective ribosome footprinting (Bohlen et al., 2020, Oh, Becker et al., 2011, Wagner et al., 2020) to determine the position on endogenous mRNAs of ribosomes containing phosphorylated Ribosomal Protein S6 (RPS6) (Figure 1A). Immunoprecipitation (IP) of ribosomes with phospho-RPS6 antibody was strongly reduced when RPS6 phosphorylation was inhibited with Torin1 (Figure 1B), indicating that the IP is specific, and that in these cells RPS6 is mainly phosphorylated by S6K downstream of mTOR. Ribosome footprints from the p-RPS6 immunoprecipitation (“pS6-ribosomes”) and from total ribosomes displayed the expected triplet periodicity, enrichment in Open-Reading Frames (ORFs) and footprint length distribution of translating 80S ribosomes (Figure 1C-C’ and Suppl. Figure 1). Generally, pS6-ribosomes have a similar distribution to total ribosomes both within transcripts (Figure 1C-C’) and between different mRNAs in the transcriptome (Figure 1D). Nonetheless, pS6-ribosomes were not equally bound to all mRNAs in the transcriptome. For every individual transcript we calculated the proportion of bound ribosomes that are phosphorylated on RPS6 by dividing the number of pS6-ribosome footprints to the number of total ribosome footprints (“pS6 abundance”). A z-vs-z analysis revealed that mRNAs from 1637 genes had lower pS6 abundance than expected by chance, and 69 genes had higher pS6 abundance than expected (Figure 1E). We reasoned that some property of these 1637 transcripts might cause them to have reduced pS6 abundance. We analyzed various features and found that the transcripts with low pS6 abundance were significantly enriched for long coding sequences (Figure 1F), but not long 5’UTRs or 3’UTRs (Figure 1 G-H).

**Figure 1:**
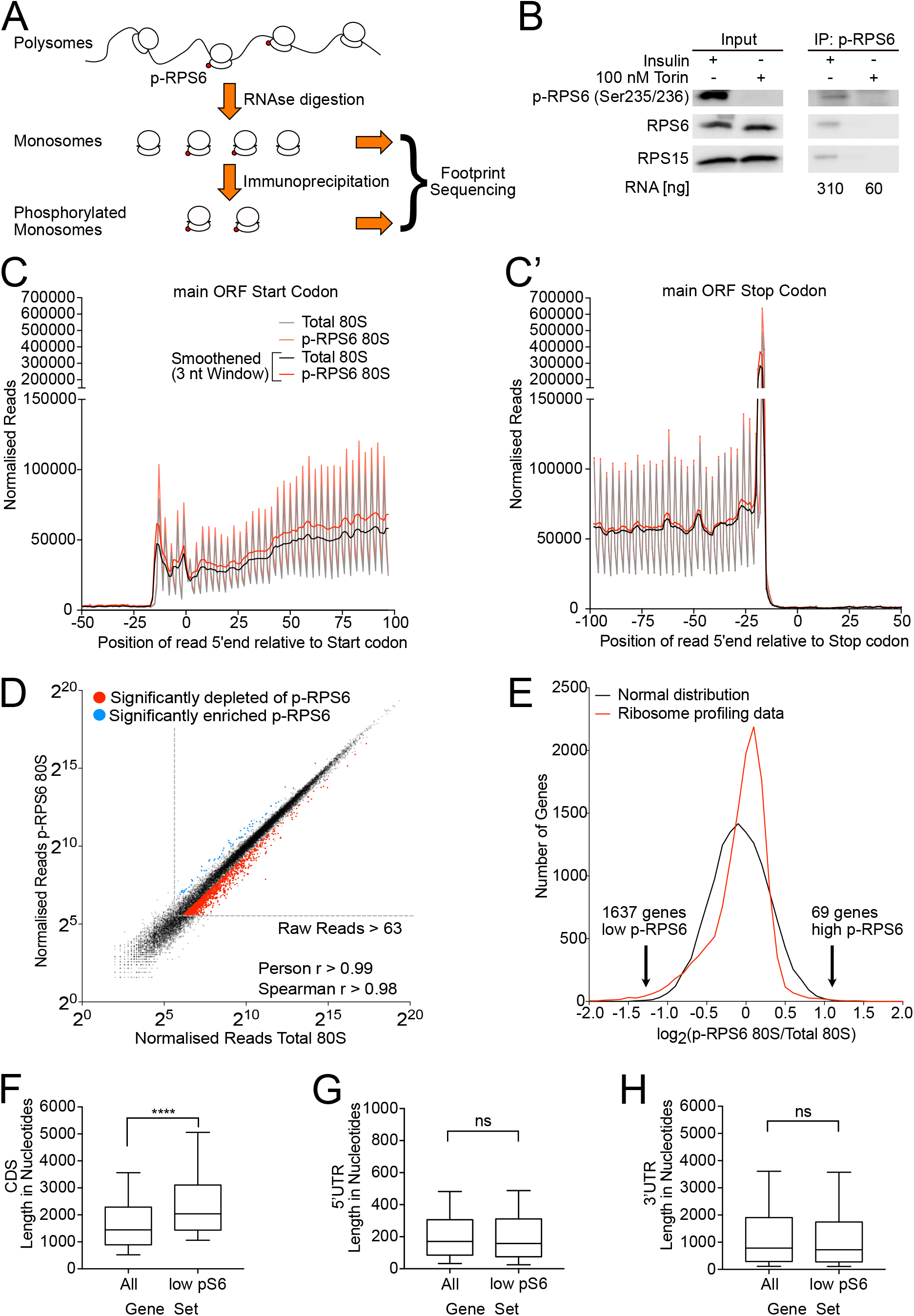
Modification-selective ribosome footprinting reveals the distribution of ribosomes containing phosphorylated RPS6 on mRNAs in vivo. **(A)** Schematic diagram illustrating phospho-RPS6-selective ribosome footprinting. Polysomes are digested by RNAse 1, monosomes are isolated by sucrose density centrifugation, ribosomes containing phosphorylated RPS6 are purified by immunoprecipitation, and footprints are then extracted and sequenced. **(B)** Verification of specificity of the p-RPS6 immunoprecipitation. Ribosomes phosphorylated on RPS6 Serine 235 and 236 were immunoprecipitated from whole cell lysates of control cells or cells treated with Torin (100 nM, 30 min). No ribosomal proteins and only background levels of RNA were obtained in the IP from Torin-treated cells. **(C-C’)** Metagene profiles for total and p-RPS6 selective 80S ribosome footprints from HeLa cells showing the position of the 5’end of ribosome footprints relative to (C) start and (C’) stop codons of all transcripts. Read counts were normalized to sequencing depth. “Smoothened” indicates the curve was smoothened with a 3nt sliding window. **(D)** Generally, p-RPS6 ribosomes are present on all mRNAs, as can be seen from the tight correlation between pRPS6-ribosome footprints and total 80S footprints. Nonetheless, some transcripts have lower (red) or higher (blue) p-RPS6 ribosome occupancy than expected (determined by z-vs-z analysis in panel E). Reads counts per gene are normalized to library sequencing depth. A minimum threshold of >63 raw reads was set. **(E)** Identification of transcripts with low or high p-RPS6 ribosome occupancy by z-vs-z analysis. Histogram of relative p-RPS6 ribosome occupancy per transcript (red) versus a normal distribution with the same mean, standard deviation, and area under the curve (black). 1637 genes with low p-RPS6 ribosome occupancy and 69 genes with high p-RPS6 occupancy were identified by z-vs-z analysis. **(F-H)** Transcripts with low p-RPS6 occupancy (n=1637) have long coding sequences (F) but not long 5’UTRs (G) or 3’UTRs (H). Line=median, boxes=upper and lower quartile boundaries, whiskers=first and last decile boundaries. P-values were determined using unpaired, two-sided Mann-Whitney tests, ****p<0.0001

### RPS6 phosphorylation is progressively removed from translating ribosomes, particularly on long open-reading frames

We sought to understand why RPS6 phosphorylation is less present on ribosomes translating long ORFs compared to short ORFs. We noticed that on long ORFs, pS6 footprints are not equally depleted across the entire length of the transcript. Instead, they are present at higher levels at the 5’ end of the ORF and then decrease, as can be seen for instance on the POLR2A mRNA (Figure 2A). This trend can be seen globally transcriptome-wide, for instance on a metagene plot of all ORFs longer than 3000nt (Figure 2B). Since this decrease occurs slowly and progressively with distance from the translation start codon, it is not visible on short ORFs, and it causes longer ORFs to have a lower average pS6 abundance when integrated over the entire transcript length. Consequently, transcriptome-wide, the longer the ORF, the lower the average pS6 abundance (Figure 2C). Since we are calculating the ratio of pS6-ribosomes to total ribosomes, this implies that the abundance of non-phosphorylated ribosomes progressively increases on long ORFs. As it appears unlikely that pS6-ribosomes are being replaced with non-phosphorylated ribosomes in the middle of peptide chain elongation, this implies that ribosomes become progressively dephosphorylated on RPS6 as they elongate. A previous report showed that S6K, the kinase responsible for phosphorylating RPS6, is physically associated to preinitiation complexes, which are 40S ribosomes bound to translation initiation factors (eIFs), and that this association occurs via eIF3 (Holz, Ballif et al., 2005). We previously showed that in human cells 40S ribosomes remain bound to eIFs while they scan 5’UTRs up to the main ORF start codon, and that indeed 80S ribosomes remain bound to eIF3 for circa 12 rounds of elongation (Bohlen et al., 2020). Hence a model consistent with all these data is that 40S subunits become phosphorylated on RPS6 by S6K during 5’UTR scanning (Figure 2D). Once the 40S converts to an 80S ribosome and undergoes circa 12 rounds of peptide elongation, eIF3 and S6K are released, and S6 becomes progressively dephosphorylated by cytosolic PP1 and / or PP2B (Andres, Johansen et al., 1987) (Figure 2D).

**Figure 2:**
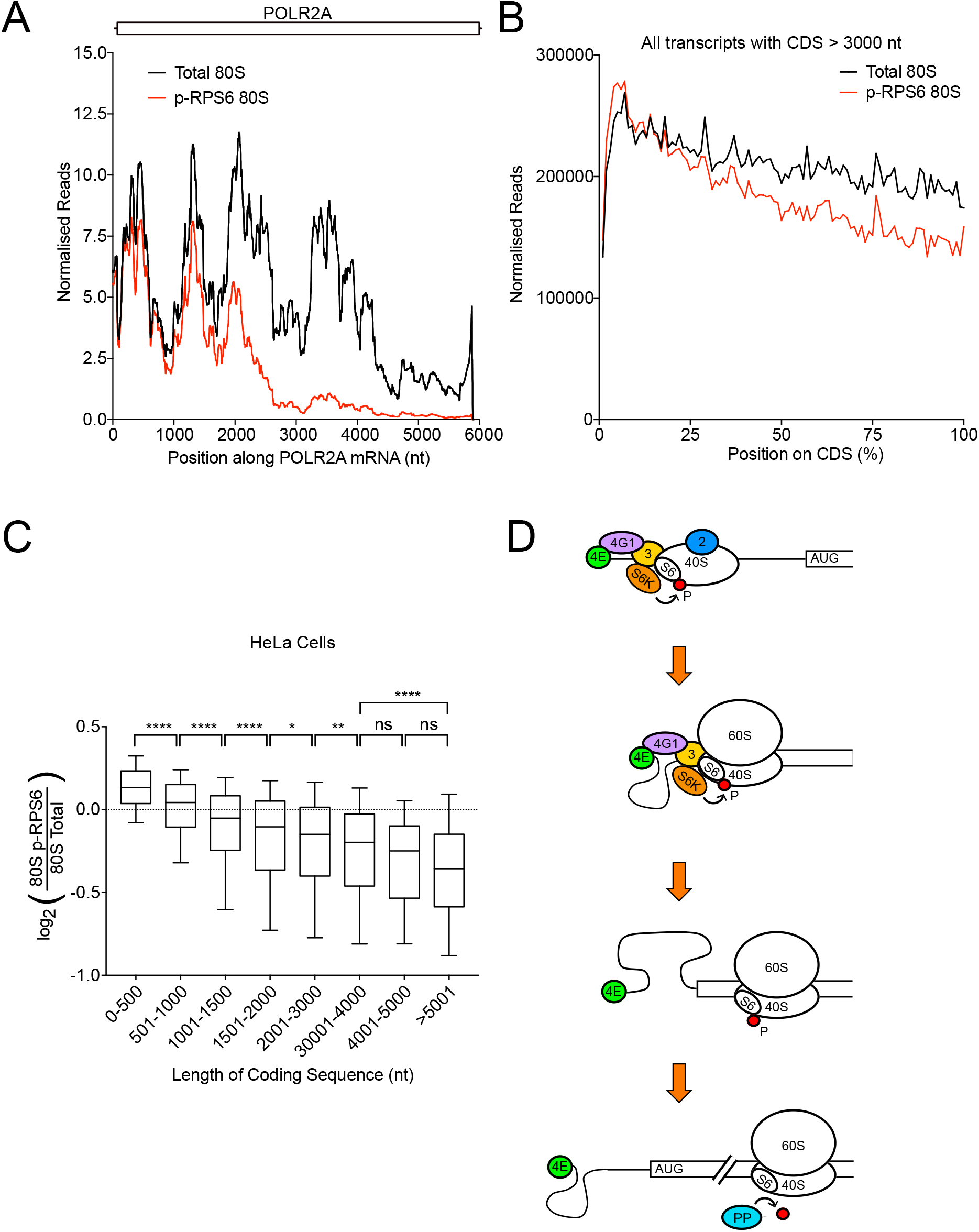
Ribosomes translating long coding sequences are progressively dephosphorylated on RPS6. **(A)** Ribosomes translating the POLR2A mRNA are progressively dephosphorylated on RPS6 Serine 235 and 236. Ribosome occupancy of total 80S and p-RPS6 80S ribosomes along the POLR2A mRNA. Curves are normalized to sequencing depth and smoothened with a sliding window of 200 nucleotides. **(B)** Progressive loss of RPS6 phosphorylation on ribosomes translating long coding sequences can be observed transcriptome-wide. Metagene plot of total 80S and p-RPS6 80S footprints on CDSs longer than 3000nt (n=4573). Position along CDS is scaled from 0% (start codon) to 100% (stop codon). Curves are normalized to sequencing depth. **(C)** Coding sequence length inversely correlates with average p-RPS6 occupancy on transcripts. Grouped analysis of the relationship between coding sequence length and p-RPS6 occupancy in HeLa cells. Transcripts were classified into the indicated groups depending on their annotated coding sequence length. Line=median, boxes=upper and lower quartile boundaries, whiskers=first and last decile boundaries. P-values were determined using unpaired, two-sided Kruksal Wallis tests, p-values were adjusted for multiple testing using statistical hypothesis testing, *p<0.0332, **p<0.0021, ****p<0.0001 **(D)** Schematic model of RPS6 phosphorylation by S6K during translation initiation and subsequent dephosphorylation during translation elongation.

### Dephosphorylation rate of RPS6 depends on mRNA environment

Although there is a general trend of lower RPS6 phosphorylation on mRNAs with longer CDS, nonetheless a scatter plot of phospho-RPS6 abundance versus CDS length shows quite a lot of dispersion (Suppl. Figure 2A), indicating that other factors also influence RPS6 abundance. To find such other factors, we did two analyses: First, we ranked all mRNAs according to their relative abundance of phospho-RPS6 reads and performed a gene set enrichment analysis using PANTHER. mRNAs coding for plasma membrane proteins, which are translated on the endoplasmic reticulum (ER), were enriched for low phospho-RPS6, whereas mRNA encoding for nuclear or cytosolic mRNAs were enriched for high phospho-RPS6 (Figure 3A, Supplemental Table 1). Secondly, we did a GO enrichment analysis on the set of mRNAs with lowest phospho-RPS6 abundance and also found an enrichment for ER-translated mRNAs and a de-enrichment for mRNAs encoding nuclear and cytosolic components (Figure 3B, Supplemental Table 2). We therefore selected all mRNAs that are translated on the ER, encoding secreted proteins, plasma-membrane integral proteins and ER/Golgi proteins based on (Jan, Williams et al., 2014) (see Materials & Methods) and found that indeed for almost all CDS length categories these mRNAs have lower phospho-RPS6 counts than average (Figure 3C-D). A position-resolved metagene plot of phospho-RPS6 abundance for ER-translated mRNAs versus all other mRNAs (Figure 3E) revealed that ribosomes translating on the ER are more rapidly dephosphorylated from nt 120 on the CDS (once the nascent signal peptide has emerged from the ribosome exit tunnel and is recognized by the Signal Recognition Particle to translocate the ribosome to the ER) to nt 400 on the CDS. After that, they are dephosphorylated with the same rate as all other mRNAs. Hence RPS6 is more rapidly dephosphorylated during early events that happen after translocation of ribosomes to the ER. In contrast, ribosomes translating mRNAs encoding nuclear proteins (GO:0031981) are more phosphorylated than average, although the magnitude of this effect is small (Figures 3F-G). Correspondingly, ribosomes translating mRNAs with long CDSs that encode for nuclear proteins tend to be more resistant to dephosphorylation as they move down the CDS (Suppl. Fig. 2B). Together, these data suggest that the rate of dephosphorylation of RPS6 depends on the subcellular environment in which the ribosomes reside.

**Figure 3:**
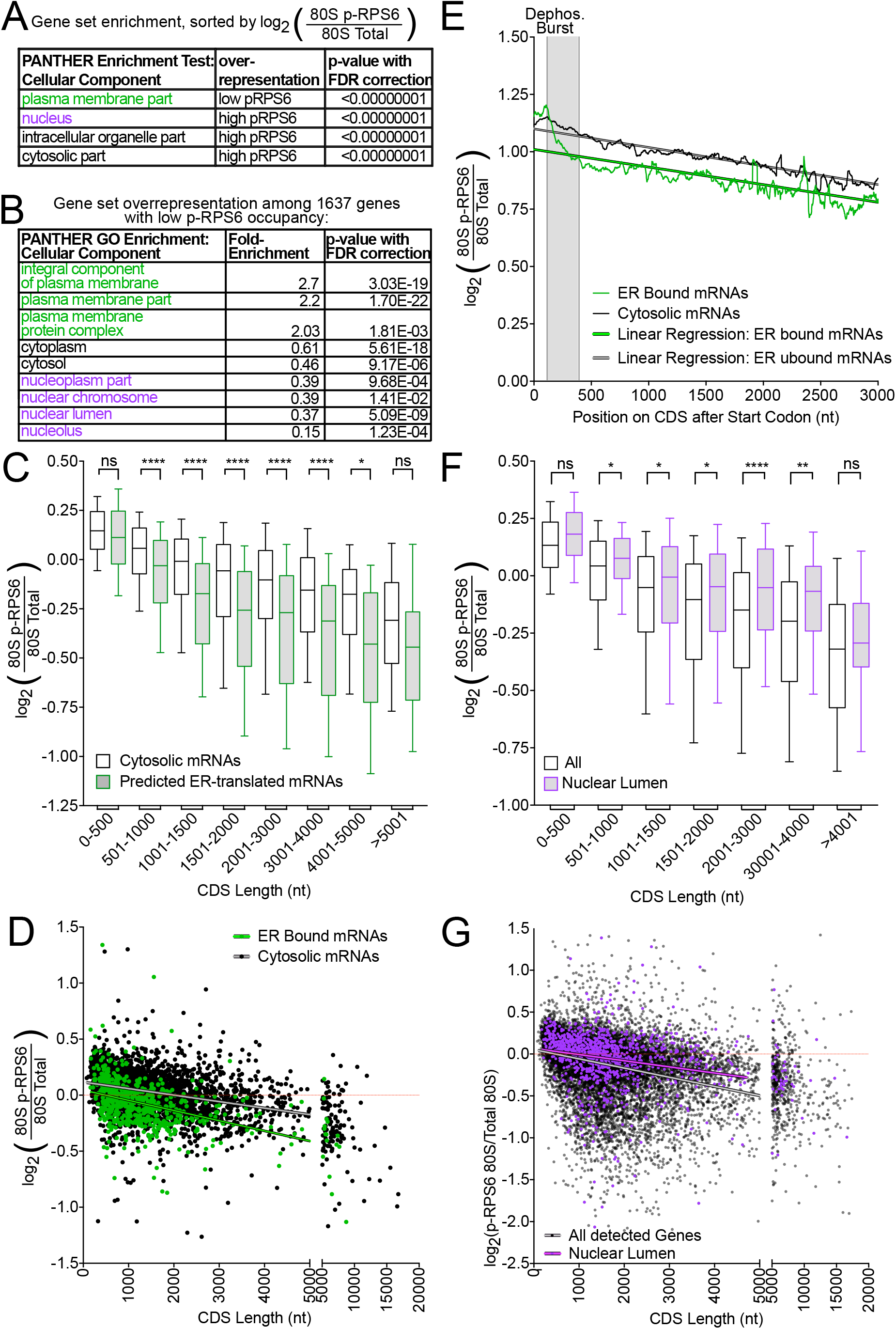
Dephosphorylation of ribosomes on RPS6 is dependent on the subcellular localization of mRNA translation. **(A-B)** Ribosomes translating mRNAs encoding plasma membrane proteins are enriched for low pRPS6 abundance while ribosomes translating mRNAs encoding nuclear components are enriched for high p-S6 abundance. (A) For each transcript the ratio of p-RPS6 / Total ribosome footprints was calculated and GO Enrichment analysis for cellular component was then computed using PANTHER. Representative, significantly enriched or depleted GO terms are shown here; The full list is provided as Suppl. Table 1 (B) Over or underrepresentation of cellular component GO terms in the 1637 genes with significantly low p-RPS6 occupancy was determined using PANTHER overrepresentation analysis. Representative, significantly enriched or depleted GO terms are shown here; The full list is provided as Suppl. Table 2. **(C-E)** Ribosomes translating secreted proteins are more strongly dephosphorylated on RPS6. (C) Grouped analysis of the relationship between coding sequence length and p-RPS6 occupancy for mRNAs predicted to be translated on the ER versus all other mRNAs. Transcripts were classified into the indicated groups depending on their annotated coding sequence length and whether they were classified as ER-associated in (Jan et al., 2014) (see Materials & Methods). Line=median, boxes=upper and lower quartile boundaries, whiskers=first and last decile boundaries. P-values were determined using unpaired, two-sided Kruksal Wallis tests, p-values were adjusted for multiple testing using statistical hypothesis testing, *p<0.0332, ****p<0.0001. (D) Scatter plot of log2(p-RPS6 Enrichment) versus coding sequence length for mRNAs experimentally validated in (Jan et al., 2014) to be ER-translated (green) or cytosolically translated (black). Linear regression of each group is also shown. (E) Metagene plot of the ratio of p-RPS6 80S to total 80S footprints on CDSs for mRNAs as in panel D. Curves are normalized to sequencing depth and smoothened with a sliding window of 40 nucleotides. Linear regression of each curve is also shown. **(F-G)** Ribosomes translating nuclear genes are more resistant to dephosphorylation on RPS6. (F) Grouped analysis of the relationship between coding sequence length and p-RPS6 occupancy for mRNAs encoding nuclear lumen proteins (GO:0031981) versus all detected genes. Transcripts were classified into the indicated groups depending on their annotated coding sequence length. Line=median, boxes=upper and lower quartile boundaries, whiskers=first and last decile boundaries. P-values were determined using unpaired, two-sided Mann Whitney tests, p-values were adjusted for multiple testing using statistical hypothesis testing, *p<0.05, **p<0.005, ****p<0.00005. (G) Scatter plot of log2(p-RPS6 Enrichment) versus coding sequence length for mRNAs encoding nuclear lumen proteins versus all other mRNAs. Linear regression of each group is also shown.

### Transcripts with short CDSs are translated less efficiently in rpS6^P−/−^ cells

We next sought to understand the functional consequences of S6 phosphorylation on translation, keeping in mind that the degree of S6 phosphorylation depends on the length of the CDS being translated. To this end, we performed pS6-selective and standard ribosome footprinting in wildtype and rpS6^P−/−^ mouse embryonic fibroblasts (MEFs), where the phosphorylated serines (235, 236, 240, 244, and 247) on endogenous rpS6 were mutated to alanine (Ruvinsky et al., 2005) (Suppl. Figure 3A-B). As in HeLa cells, the ribosome footprinting in MEFs yielded the expected triplet periodicity and enrichment in CDSs of translating 80S ribosomes (Suppl. Figure 3C-C’). As expected, the mRNAs with lowest pS6-abundance transcriptome-wide were mitochondrially encoded mRNAs (Suppl. Figure 3D). (We did not detect mitochondrial mRNAs in our HeLa footprinting experiments.) As in human cells, mRNAs translated on the ER were enriched among the transcripts with low pS6 abundance, whereas cytosolically-translated mRNAs were de-enriched (Suppl. Table 3). Likewise, we observed reduced levels of pS6 on long transcripts in MEFs (Figure 4A), although the effect was weaker than in HeLa cells. Nonetheless, the progressive loss of RPS6 phosphorylation is clearly visible on single transcripts with long CDSs such as EP400 (Figure 4B) and on a metagene plot for all transcripts with long CDS >3000 nt (red trace, Figure 4D), indicating it is a conserved feature of translation in humans and mice. Interestingly, when comparing the translation efficiency for each transcript in rpS6^P−/−^ versus wildtype MEFs, we found that transcripts with short CDSs showed reduced translational efficiency in rpS6^P−/−^ MEFs in comparison to transcripts with longer CDSs (Figure 4C). This effect is mostly driven by changes in the number of ribosome footprints per transcript (Suppl. Figure 3E) and not by changes in mRNA levels (Suppl. Figure 3F), indicating that a translational mechanism is at play. This indicates that in wildtype cells the high levels of RPS6 phosphorylation on transcripts with short CDSs helps promote their translation in comparison to transcripts with longer CDSs where RPS6 phosphorylation is depleted.

**Figure 4:**
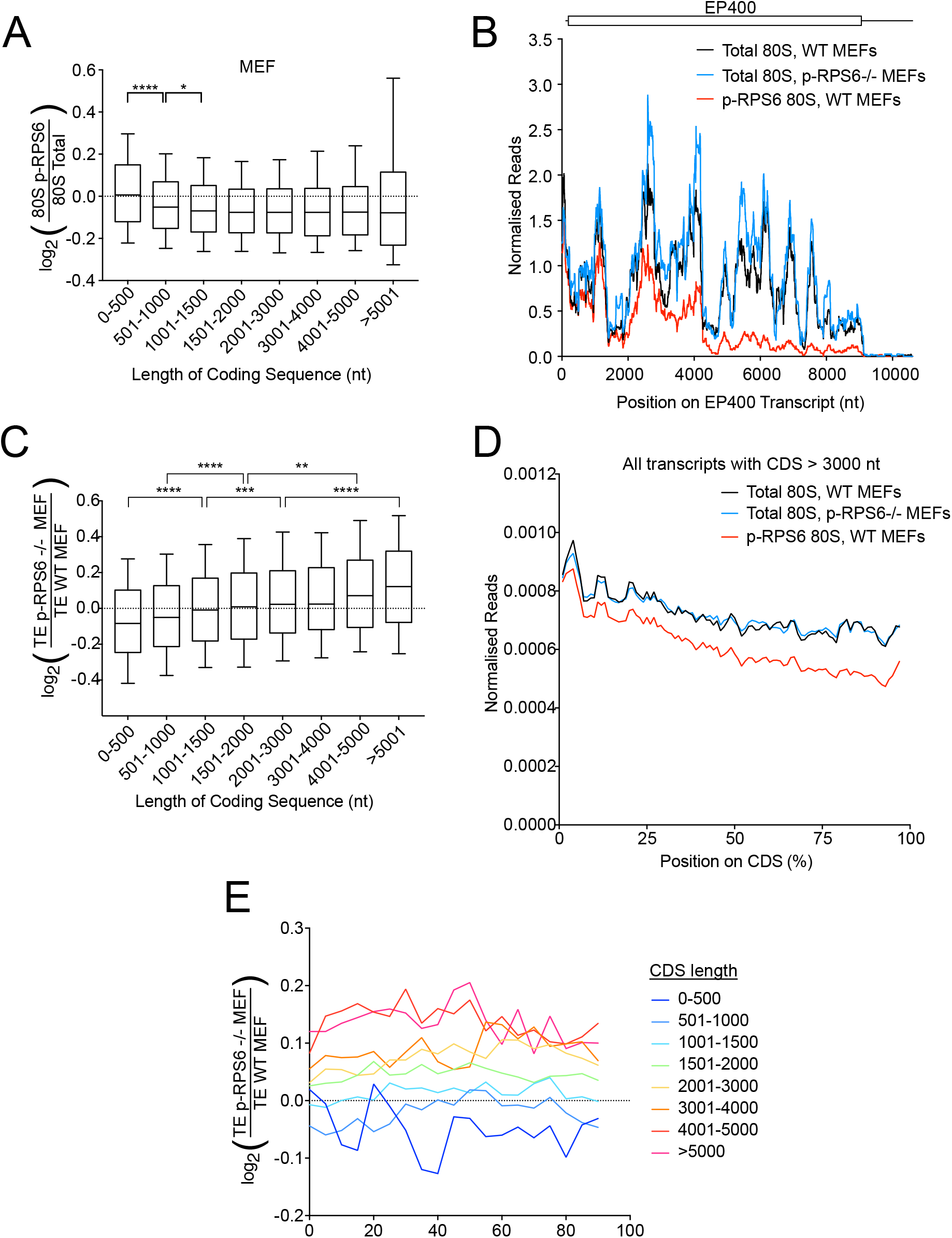
Phosphorylation of RPS6 promotes translation of transcripts with short CDSs. **(A)** As in human cells, average occupancy of phospho-RPS6 ribosomes significantly drops in transcripts with longer CDSs, although the effect is not as large as in HeLa cells. Grouped analysis of the relationship between coding sequence length and p-RPS6 occupancy in mouse embryonic fibroblasts. Transcripts were classified into the indicated groups depending on their annotated coding sequence length. Line=median, boxes=upper and lower quartile boundaries, whiskers=first and last decile boundaries. P-values were determined using unpaired, two-sided Kruksal Wallis tests, p-values were adjusted for multiple testing using statistical hypothesis testing, *p<0.0332, ****p<0.0001 **(B)** Ribosomes translating the EP400 mRNA are progressively dephosphorylated on RPS6 Serine 235 and 236. Ribosome occupancy of total 80S and p-RPS6 80S ribosomes along the EP400 mRNA. Curves are normalized to sequencing depth and smoothened with a sliding window of 200 nucleotides. **(C)** Short transcripts are poorly translated in rpS6^P−/−^ MEFs compared to wildtype controls. Grouped analysis of the relationship between coding sequence length and relative translational efficiency in rpS6^P−/−^ MEFs compared to control MEFs. Genes were classified into the indicated groups depending on their annotated coding sequence length. Line=median, boxes=upper and lower quartile boundaries, whiskers=first and last decile boundaries. P-values were determined using unpaired, two-sided Kruksal Wallis tests, p-values were adjusted for multiple testing using statistical hypothesis testing, **p<0.0021, ***p<0.0002, ****p<0.0001 **(D)** Progressive dephosphorylation of RPS6 on ribosomes translating long coding sequences in MEFs. Metagene plot of footprints on CDSs longer than 3000nt (n=5342). Position along CDS is scaled from 0% (start codon) to 100% (stop codon). Curves normalized to sequencing depth. **(E)** Ribosome occupancy is uniformly affected across the entire length of the CDS depending on CDS length in rpS6^P−/−^ MEFs. Transcripts were classified into the indicated groups depending on their annotated coding sequence length. Curves denote the ratio in ribosome occupancy between rpS6^P−/−^ MEFs and control MEFs at various positions along the coding sequence, scaled from 0% (start codon) to 100% (stop codon). Curves normalized to sequencing depth.

Since RPS6 becomes progressively dephosphorylated on long CDSs as ribosomes move down the CDS, we asked whether we could observe position-dependent effects on translation in the rpS6^P−/−^ MEFs. In theory, one could expect that loss of RPS6 phosphorylation at the 5’end of the CDSs, where RPS6 phosphorylation is normally high, could lead to a change in ribosome density, whereas at the 3’end where RPS6 levels are low in wildtype MEFs, loss of S6 phosphorylation would have no effect. This, however, was not the case; Ribosome footprint density was similar along the entire length of long CDSs in WT and rpS6^P−/−^ MEFs (Figure 4D). Therefore, position-dependent differences in pS6 abundance do not lead directly to position-dependent changes in ribosome density.

Interestingly, in both wildtype and rpS6^P−/−^ MEFs (as well as in HeLa cells), total 80S ribosome density decreases with distance from the 5’end of the CDSs (black curves in Fig. 2D & 4D). Assuming that few ribosomes abort protein synthesis prematurely, this suggests that ribosomes progressively speed up (analogous to cars speeding up after a traffic jam, leading to lower car density). This, however, occurs to the same extent in both WT and rpS6^P−/−^ MEFs (Figure 4D). Hence, elongation speed does not appear to be affected by RPS6 phosphorylation. Furthermore, transcripts with long CDSs had uniformly elevated ribosome density in rpS6^P−/−^MEFs throughout the entire length of the CDS, from the start codon onwards (Figure 4E). Thus, through an unknown mechanism which will require further investigation, the phosphorylation of RPS6 on ribosomes when they terminate translation of short CDSs seems to increase initiation rates on that transcript. One can speculate this occurs due to circularization of translated mRNAs.

### Translation of 5’ TOP mRNAs is resistant to loss of RpS6 phosphorylation

We noticed that one set of transcripts appears to defy the global trends described above. As discussed above, transcripts that are poorly translated in Rps6^P−/−^ MEFs are the ones with short CDSs (Figure 4C). We asked what transcripts are present in the opposite category – the ones that retain efficient translation in rpS6^P−/−^ MEFs. In addition to the expected transcripts with long CDSs (Figure 4C) we also found mRNAs containing 5’ terminal oligopyrimidine tract (TOP) motifs: A Gene Ontology (GO) enrichment analysis on the set of transcripts with a high ratio of translation efficiency in rpS6^P−/−^ MEFs versus wildtype MEFs found transcripts coding for ribosomal proteins to be most highly enriched (Figure 5A). Indeed, when comparing rpS6^P−/−^ to wildtype MEFs, ribosomal proteins are even better translated on average than transcripts with CDSs longer than 5000 nucleotides (Figure 5B, Suppl. Figure 4A). This was surprising because ribosomal proteins are short (median CDS length of 455 nt) and therefore would be expected to be translated poorly in rpS6^P−/−^ MEFs, like other short ORF mRNAs (Figure 4C). Transcripts encoding ribosomal proteins contain 5’ TOP motifs (Cockman, Anderson et al., 2020). We found that other TOP mRNAs that do not code for ribosomal proteins were also translated more efficiently in rpS6^P−/−^ MEFs compared to transcripts with CDSs of similar length (Figure 5B, Suppl. Figure 4A). One explanation could be that, despite their short length, TOP mRNAs are preferentially translated by ribosomes lacking RpS6 phosphorylation, and hence are insensitive to loss of RpS6 phosphorylation. This, however, was not the case: In both HeLa and MEFs, ribosomal and TOP mRNAs have high levels of p-RPS6-ribosome footprints (Figure 5C-D, Suppl. Figure 4B-D). Hence TOP-motif-containing mRNAs continue being efficiently translated in rpS6^P−/−^ MEFs despite losing phosphorylation on RpS6, suggesting they are translated using a mechanism that bypasses the requirement for RpS6 phosphorylation, perhaps due to a role of LARP1 (Cockman et al., 2020). This may provide a hint to the mechanism how RpS6 phosphorylation affects translation initiation. In agreement with these footprinting data, rpS6^P−/−^ MEFs have elevated levels of ribosomal proteins compared to control MEFs (Figure 5E-F). This is not due to a general increase in mTOR signaling as a compensation to p-RPS6 mutation since mTOR target phosphorylation is unchanged in rpS6^P−/−^ MEFs (Suppl. Figure 4E). Increased levels of ribosomes may explain the global increase in protein synthesis rates previously observed in rpS6^P−/−^ MEFs (Ruvinsky et al., 2005).

**Figure 5:**
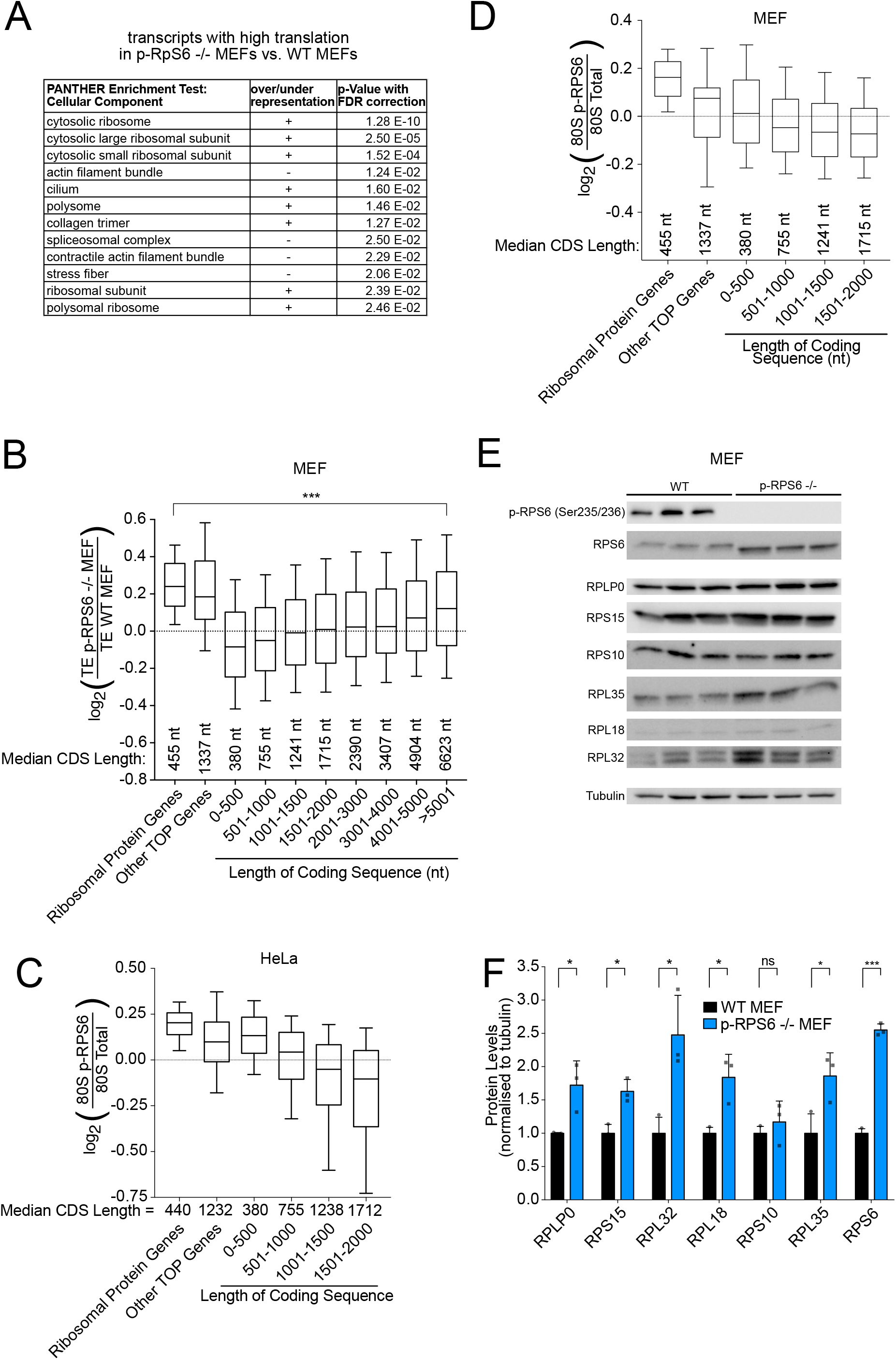
TOP mRNAs are efficiently translated independent of RPS6 phosphorylation. **(A)** mRNAs encoding ribosomal proteins are enriched among the set of transcripts whose translation efficiency drops least in rpS6^P−/−^ MEFs compared to control MEFs. For each transcript the ratio of (translation efficiency in rpS6^P−/−^ MEFs) / (translation efficiency in control MEFs) was calculated and GO Enrichment analysis for cellular component was then computed using PANTHER. **(B)** Ribosomal protein and TOP-containing mRNAs escape the translational repression observed for mRNAs with short CDSs in rpS6^P−/−^ MEFs. Grouped analysis of translational efficiency in mouse embryonic fibroblasts. Genes were classified into the indicated groups. Line=median, boxes=upper and lower quartile boundaries, whiskers=first and last decile boundaries. P-values were determined using unpaired, two-sided Kruksal Wallis tests, p-values were adjusted for multiple testing using statistical hypothesis testing, ***p<0.0002 **(C-D)** Ribosomal protein and TOP-containing mRNAs are translated by ribosomes phosphorylated on RPS6 in both HeLa (C) and MEF cells (D). Grouped analysis of p-RPS6 occupancy for transcripts grouped as indicated. Line=median, boxes=upper and lower quartile boundaries, whiskers=first and last decile boundaries. **(E-F)** rpS6^P−/−^ MEF cells have elevated levels of ribosomal proteins. (E) Western blot analysis of ribosomal protein levels in wildtype control and rpS6^P−/−^ MEF cells. (F) Three biological replicates of whole cell lysates from each genotype were normalized to each other for total protein and subjected to immunoblot analysis. Tubulin was used as a loading control. (E) Quantification of protein band intensity normalized to tubulin. P-values were calculated using unpaired, two-sided t-tests. *p< 0.05, ***p< 0.0005

## DISCUSSION

We previously developed a method for selective 40S and 80S ribosome profiling in human cells which employs an immunoprecipitation strategy to identify the location on endogenous mRNAs of ribosomes bound to a factor of interest, such as an initiation factor (Bohlen et al., 2020). Here we extend that method to look at ribosomes containing a post-translational modification of interest, in this case phosphorylation of RPS6. This modification-selective ribosome footprinting is a method that can be used in the future to study also other ribosome modifications, such as ribosome ubiquitination upon UPR activation (Higgins et al., 2015), ribosome N(*α*)-acetylation which promotes ribosome activity (Kamita et al., 2011) as well as modifications of ribosome interacting proteins such as translation initiation factors (Duncan & Hershey, 1984, Morley, Coldwell et al., 2005, Proud, 2015).

We find that RPS6 phosphorylation is highest on ribosomes positioned on the start codons of main Open Reading Frames (ORF), and then progressively decreases as ribosomes translate the ORF, making this drop most relevant on mRNAs with long ORFs. It was previously shown that S6K, the kinase that phosphorylates RPS6, is physically associated to pre-initiation complexes – scanning 40S ribosomes that are bound to initiation factors – via eIF3 (Holz et al., 2005). We previously showed that ribosomes are bound to eIF3 during the entirety of the scanning process and during the first few rounds of peptide elongation, after which they let go of eIF3 (Bohlen et al., 2020). Hence one likely molecular model that could explain these observations is that RPS6 becomes phosphorylated by S6K while it is part of a scanning 40S ribosome that is associated to eIF3 and hence S6K. After the first few rounds of peptide elongation, the 80S ribosome lets go of eIF3, and hence S6K, thereby allowing for the phosphatase-mediated progressive dephosphorylation of RPS6 (Fig. 2D).

Interestingly, we find that the rate of dephosphorylation of RPS6 as a ribosome translates a CDS is dependent on the subcellular localization of the ribosome. Ribosomes translating mRNAs on the ER have an initial phase of dephosphorylation that is more rapid that that of ribosomes in the cytosol. This suggests the local milieu affects the dephosphorylation rate. Given that ribosomes translating mRNA encoding nuclear proteins show the opposite trend – elevated RPS6 phosphorylation – this suggests these mRNAs may also reside in a particular subcellular location or milieu. Although we describe here two factors that explain variation in pS6 abundance (CDS length and mRNA localization) additional factors likely also influence this to explain all of the observed variation (Suppl. Fig. 2A).

The data we provide here indicate that RPS6 phosphorylation has a functional consequence on translation. In cells lacking RPS6 phosphorylation we observe a drop in translation efficiency of mRNAs with short CDSs, which normally have high RPS6 phosphorylation. In contrast, mRNAs with longer CDSs, and hence lower RPS6 phosphorylation, do not show a drop in translation efficiency when RPS6 phosphorylation is eliminated in rpS6^P−/−^ mutant MEFs. Mysteriously, the state of RPS6 phosphorylation of ribosomes as they reach the end of a CDS seems to affect translation efficiency on the entirety of that mRNA – i.e. initiation rates on an mRNA are increased if ribosomes terminating translation on that mRNA are still phosphorylated on RPS6. We do not know how this 3’-to-5’ information flow works mechanistically, however previous studies suggest such a flow exists (Fernandes, Moura et al., 2017). This could occur if mRNAs are circularized via interactions between the poly-A tail and the 5’cap and RPS6 phosphorylation promotes ribosome re-loading for a new round of translation. On longer ORFs, ribosomes would be preferentially dephosphorylated when they terminate, reducing initiation rates. Since all ribosomes are un-phosphorylated in rpS6^P−/−^ cells, this relative disadvantage of long ORFs would disappear, explaining the observed de-repression. Alternatively, RPS6 could affect circularization itself. A hint to the relevant mechanism comes from the fact that mRNAs containing 5’ TOP motifs seem to evade this regulatory mechanism. Further work will be necessary to unravel these mechanisms.

Interestingly, we noticed that on mRNAs with longer CDSs ribosome density decreases towards the 3’ end of CDSs transcriptome-wide both in human cells (Figure 2B) and in mouse cells (Fig. 4D). This indicates that ribosomes progressively increase their translation speed towards the 3’ end of the CDS. This could potentially prevent ribosome collisions, which would occur if ribosomes translate with a uniform speed. As ribosomes translate, stochastic events cumulatively influence the distance separating two ribosomes, such that a distance that is sufficient to prevent collisions early at the 5’ end of an ORF may become insufficient after many rounds of elongation (Suppl. Figure 5). A progressive acceleration in translation speed could counteract this effect. We speculated that RPS6 phosphorylation might be mediating this effect. This, however, does not appear to be the case. Ribosome density decreases equally towards CDS 3’ends in control and rpS6^P−/−^ MEFs (Fig. 4D), indicating that ribosomes accelerate both in the presence and absence of RPS6 phosphorylation. An alternate interpretation to the drop in ribosome density is ribosome drop-off during translation (Sin, Chiarugi et al., 2016), however from our graph (Figs. 2B & 4D) this would indicate that ∼30% of all large proteins are synthesized as truncated proteins.

In sum, we identify here a position-dependent effect on ribosomal phosphorylation. Future work will hopefully shed light on how ribosome phosphorylation at the end of a CDS can affect initiation rates on that mRNA.

## MATERIALS & METHODS

### Cell Culture

HeLa cells and MEF cells were cultured in DMEM +10% fetal bovine serum +100 U/ml Penicillin/Streptomycin (Gibco 15140122). Cells were sub-cultured using Trypsin-EDTA for dissociation.

### Immunoblotting

Cells were lysed using standard RIPA lysis buffer (150 mM NaCl, 1% NP40, 1% (w/v) Sodium Deoxycholate, 0.1% SDS, 50 mM Tris pH 8) containing protease inhibitors (Roche mini EDTA-free, 1 tablet in 10 ml) and phosphatase inhibitors (2mM Sodium Ortho-Vanadate, Roche Phosstop 1 tablet in 10 ml, 0.1 M Sodium Fluoride, 0.1 M beta-Glycerophosphate) and Benzonase (50 U/ml), after washing briefly with FBS-free DMEM. Lysates were clarified and protein concentration was determined using a BCA assay. Equal protein amounts were run on SDS-PAGE gels and transferred to a nitrocellulose membrane with 0.2 µm pore size. After Ponceau staining, membranes were incubated in 5% skim milk in PBST (134 mM NaCl, 2,7 mM KCl, 5,4 mM Na_2_HPO_4_, 1.47 mM KH_2_PO_4_) for 1 hour, briefly rinsed with PBST and then incubated in primary antibody solution (5% BSA PBST or 5% skim milk PBST) overnight at 4C. Membranes were then washed three times 15 minutes each in PBST, incubated in secondary antibody solution (1:10000 in 5% skim milk PBST) for 1 hour at room temperature, then washed again three times for 15 minutes. Finally, chemiluminescence was detected using ECL reagents and the Biorad ChemiDoc Imaging System. No membranes were stripped. Antibodies used in this study are listed in Suppl. Table 4.

### Immunoprecipitation

One million HeLa cells or 0.5 million MEF cells were seeded per well of a 6-well dish. Cells were treated with 100 nM Torin1 for 30 minutes. Cells were washed briefly with FBS-free growth medium, then lysed in 200 µl Lysis Buffer (0,5 M HEPES pH 7.5, 10 mM MgCl2, 0.2 M KCl, 1% NP40, 200 μM CHX, 0.1 M NaF, 0.011 g/ml β-Glycerophosphate, 2 mM Sodium Vanadate, PhosSTOP**^™^** (Roche 04 906 845 001) and cOmplete^™^ Mini (Roche 11836153001) both at 2x the suppliers indicated concentration, tablet dissolved in water) per well. Cells were scraped off the plate and the collected lysate was transferred to a 1.5 ml reaction tube. Lysates were clarified at 20.000g for 10 minutes, at 4°C. Input samples were saved by transferring 80 µL of the lysate into a fresh tube, and the rest of the lysate was used for immunoprecipitation: Protein A magnetic beads (Thermo 10001D) were prepared according to the manufacturer’s instructions. For the IP from HeLa cell lysates, 100 µl of magnetic beads and 10 µl of anti-p-RPS6 (Ser235/236) (Cell signaling #4857) were used per condition. For the IP from MEFs, 50 µl of magnetic beads and 20 µl of antibody were used per condition. Beads were washed three times and then added to the lysates. Lysates with beads were incubated for 2 hours, rotating at 4°C. Then beads were washed three times with bead wash buffer (20 mM Tris pH 7.4, 10 mM MgCl2, 140 mM KCl, 1% NP40), including a change of tube during the last wash. Beads were split into two aliquots. To one of them 1x Laemmli buffer was added an beads were boiled at 95°C for 5 minutes for western blot analysis. The other half was subjected to RNA extraction using TRIzol reagent (Invitrogen 15596026) according to the manufacturer’s instructions. RNA was then analysed on an Agilent Bioanalyzer using the total RNA chip.

### Immunofluorescence

MEF cells were seeded at 500.000 cells per well of a 6-well plate on microscopy coverslips. Medium was removed and cells were fixed with 4% formaldehyde in PBS for 20 minutes at room temperature. Samples were then washed three times with PBS, then blocked for 45 minutes with PBS 0.2% Triton 0.1% BSA. Primary anti-p-RPS6 (1:1000, Ser235/236, Cell signaling #4857) antibody was incubated for 2 hours in PBS, 0.2% Triton, 0.1% BSA at room temperature. Samples were then washed four times with PBS, 0.2% Triton, 0.1% BSA, and incubated for 2 hours at room temperature with secondary antibody (anti-rabbit, 1:10.000) in PBS, 0.2% Triton, 0.1% BSA. After washing four times with PBS 0.2% Trition, including DAPI in the third wash, samples were equilibrated in mounting medium for 10 minutes. Cover slips were then mounted on a slide holder and images were taken using a standard cell culture fluorescence microscope.

### p-RPS6 selective and total ribosome footprinting

HeLa cells were seeded in ten 10 cm dishes at 1.5 million cells per dish in 10 ml growth medium. MEF cells were seeded in three 15 cm dishes per condition at 3 million cells in 20 ml growth medium. Two days later, cells were harvested for Ribo-seq. Cells were briefly rinsed with ice-cold PBS containing 10 mM MgCl2, 200 μM cycloheximide (CHX). This solution was poured off, removed by gently tapping the dish onto paper towels, and cells were lysed with 200 µl of lysis buffer (0,1 M HEPES pH 7.5, 20 mM MgCl2, 0.4 M KCl, 2% NP40, 400 μM CHX, 0.2 M NaF, 0.022 g/ml β-Glycerophosphate, 4 mM Sodium Vanadate, PhosSTOP**^™^** (Roche 04 906 845 001) and cOmplete^™^ Mini (Roche 11836153001) both at 2x the suppliers indicated concentration, tablet dissolved in water) per plate. Cells were scraped off and lysate was collected in a 1.5mL tube. The collected volume was roughly 400 µl per plate. After brief vortexing, lysate was clarified by centrifuging for 10 minutes at 20.000 g at 4C. Approximate RNA concentration was measured using a Nanodrop system and 100U of Ambion RNAse 1 was added per 120 µg of measured RNA. To prepare undigested polysome profiles RNAse was omitted. Lysates were incubated with RNAse for 5 minutes on ice. Lysates were then pipetted onto 17.5-50% sucrose gradients, which were produced by freezing and layering 50% (2.5ml), 41.9% (2.5ml), 33.8% (2.5ml), 25,6% (2.5ml) and 17.5% (1.8ml) sucrose solutions (10 mM Tris HCl pH 7.4, 10 mM MgCl2, 140 mM KCl, 200 µM CHX) in Seton Scientific Polyclear Tubes 9/16×3-3/4 IN, and centrifuged at 35.000 rpm for 3.5 hours in Beckmann SW40 rotor. Gradients were fractionated using a Biocomp Gradient Profiler system and 80S fractions were collected for footprint isolation and immunoprecipitation. 500 µl of 80S fractions were saved for total footprint isolation.

For immunoprecipitation, antibodies were bound to protein A magnetic dynabeads (Thermo 10001D) according to the manufacturer’s instructions. For the IP from HeLa cells lysates, 100 µl magnetic beads and 60 µl anti-p-RPS6 (Ser235/236) (Cell signaling #4857) were used. For the preparative IP from MEFs, 180 µl magnetic beads and 100 µl antibody were used. Beads were washed three times and then added to the 80S fractions. Fractions with beads were incubated for 2 hours, rotating at 4°C. Then beads were washed three times with bead wash buffer (20 mM Tris pH 7.4, 10 mM MgCl2, 140 mM KCl, 1% NP40), including a change of tube during the last wash. Bead volume was increased to ∼500µl with bead wash buffer. 10% of the sample volume was saved for western blotting analysis of the IP. Total footprint fractions and IPed fractions were then subjected to RNA extraction: 55 µl (1/9th of volume) of 20% SDS was added, 650 µl Acid-Phenol Chloroform (Ambion) was added and the mixture was incubated at 65°C with 1300 rpm shaking for 45 minutes. Tubes were then placed on ice for 5 minutes, spun for 5 min at 20.000 g and the supernatant was washed once with acid-phenol chloroform and twice with chloroform. RNA was then precipitated with isopropanol, analyzed on an Agilent Bioanalyzer system to asses RNA integrity, and subjected to library preparation (see below). For RNA-seq in MEF cells, 3 million cells were seeded in 15 cm dishes in 20 ml growth medium and harvested two days later using TRIzol according to the manufacturer instructions.

### Deep-sequencing library preparation

For 80S footprinting and RNA-seq, libraries were prepared as follows: Samples were depleted of ribosomal RNA using the Illumina Ribo-Zero Gold kit. Poly-A mRNA was purified from total RNA using the NEB polyA Spin mRNA Isolation Kit. Poly-A mRNA was then fragmented using chemical cleavage in 50 mM NaHCO3 at pH 10, 95°C for 12 minutes. Then total RNA was processed in parallel with the depleted RNA from 80S ribosome fractions. For size selection, RNA was run on 15% Urea-Polyacrylamide gels and fragments from 25-35 nt were excised using reference ssRNA nucleotides of 25 and 35 basepairs run on a neighboring lane. RNA was extracted from the gel pieces and phosphorylated using T4 PNK. Deep sequencing libraries were prepared from these RNA fragments using the Bio-Scientific NEXTflex Small RNA-Seq Kit v3. DNA was amplified with 11 PCR cycles for the HeLa Ribo-seq samples and 9-14 cycles for the MEF samples. Deep-sequencing libraries were sequenced on the Illumina Next-Seq 550 system.

### Data analysis and statistics

Analysis of ribosome footprinting NGS data: Adapter sequences and randomized nucleotides were trimmed from raw reads using cutadapt (https://doi.org/10.14806/ej.17.1.200). Ribosomal RNA and tRNA reads were removed by alignment to human tRNA and rRNA sequences using bowtie2 (Langmead & Salzberg, 2012). Then, the remaining reads were separately aligned to the human transcriptome (Ensembl transcript assembly 94) and human genome (hg38) using BBmap (sourceforge.net/projects/bbmap/). Generally, counted reads were normalized to sequencing depth (number of alignments per library). Read counting (Figs. 1D-H, 2C, 3A, 3C, 4A-D, S1A, S2D-E, S3A-C), metagene plots (Figs. 1C-C’, 2B, 3D, 3E, S2C-C’, S3D), single transcript traces (Figs. 2A and 3B), metagene plots with position buckets (Fig. 4E), as well as removal of transcripts with PCR artefacts were done with custom software written in C available on GitHub (https://github.com/aurelioteleman/Teleman-Lab). Translation efficiency (TE) was calculated from the number of 80S ribosome footprints in a coding sequence divided by the number of RNA sequencing reads in a coding sequence. For PANTHER Gene Set Enrichment test (Fig. 4A), a list of all detected genes (raw reads >63) and their change in translation between control wildtype and rps6^p−/−^ cells was entered into the panther suite (Mi, Ebert et al., 2021) and enrichment for gene sets of cellular components was calculated. Bioinformatic classification of ER-translated versus cytosolic mRNAs (Fig. 3C) was taken from (Jan et al., 2014), as described in their Supporting Materials and Methods: “Similar to the yeast secretome, we defined the mammalian secretome as the set of proteins predicted to have a signal peptide or transmembrane domain but excluding known mitochondrial proteins. The set of proteins predicted to contain a signal peptide or transmembrane domain by Phobius were filtered to remove proteins annotated in MitoCarta (53) or associated with the gene ontology term GO:005739 (cellular component “mitochondrion”) to yield the mammalian secretome.” Classification of mRNAs experimentally determined to be translated on the ER (Fig. 3D-E) was also taken from (Jan et al., 2014): In their Supplemental Table S6, column S (log_2_(Enrichment pulldown versus input)) was used and a threshold of >0.5 was set to classify the detected mRNAs (n = 4023) into ER bound (n = 620) or unbound (n = 3403).

### Data and Software Availability

All deep sequencing datasets have been submitted to NCBI Geo (<u>GSE168977</u>). Custom software is available on GitHub: https://github.com/aurelioteleman/Teleman-Lab.

## ACKNOWLEDGEMENTS

We thank Oded Meyuhas for kindly providing the p-S6 mutant MEF cells.

## AUTHOR CONTRIBUTIONS

J.B. performed the experiments. J.B and A.T designed the work, analyzed data, interpreted data, and wrote the manuscript.

## COMPETING INTERESTS

Authors declare no competing interests.

---

**Suppl. Figure 1:**
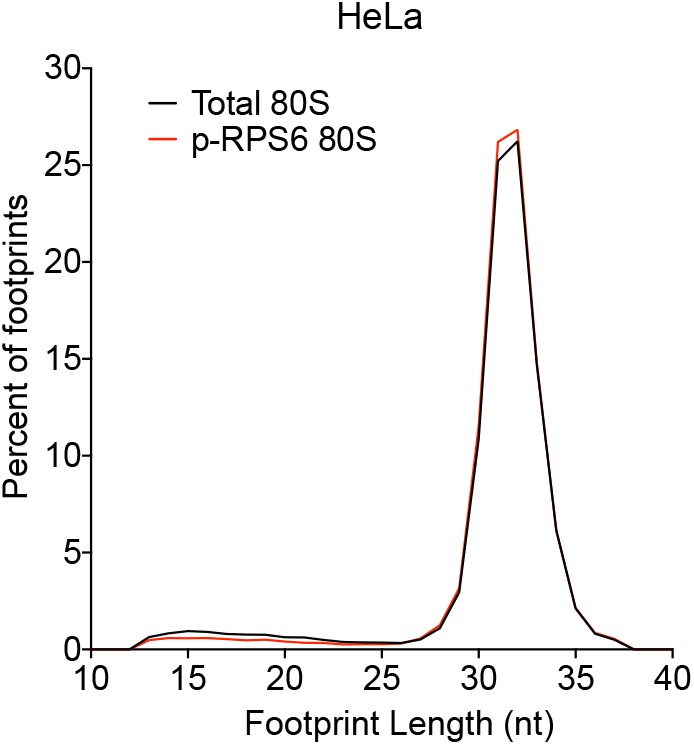
Footprint size distribution as a quality control for total 80S and p-RPS6-selective ribosome footprinting. Ribosome footprints of total 80S and p-RPS6 selective 80S from HeLa cells exhibit typical footprint lengths. Histogram of ribosome footprint length of footprints mapping into a 200 nt window around mORF start codons in the indicated samples.

**Suppl. Figure 2:**
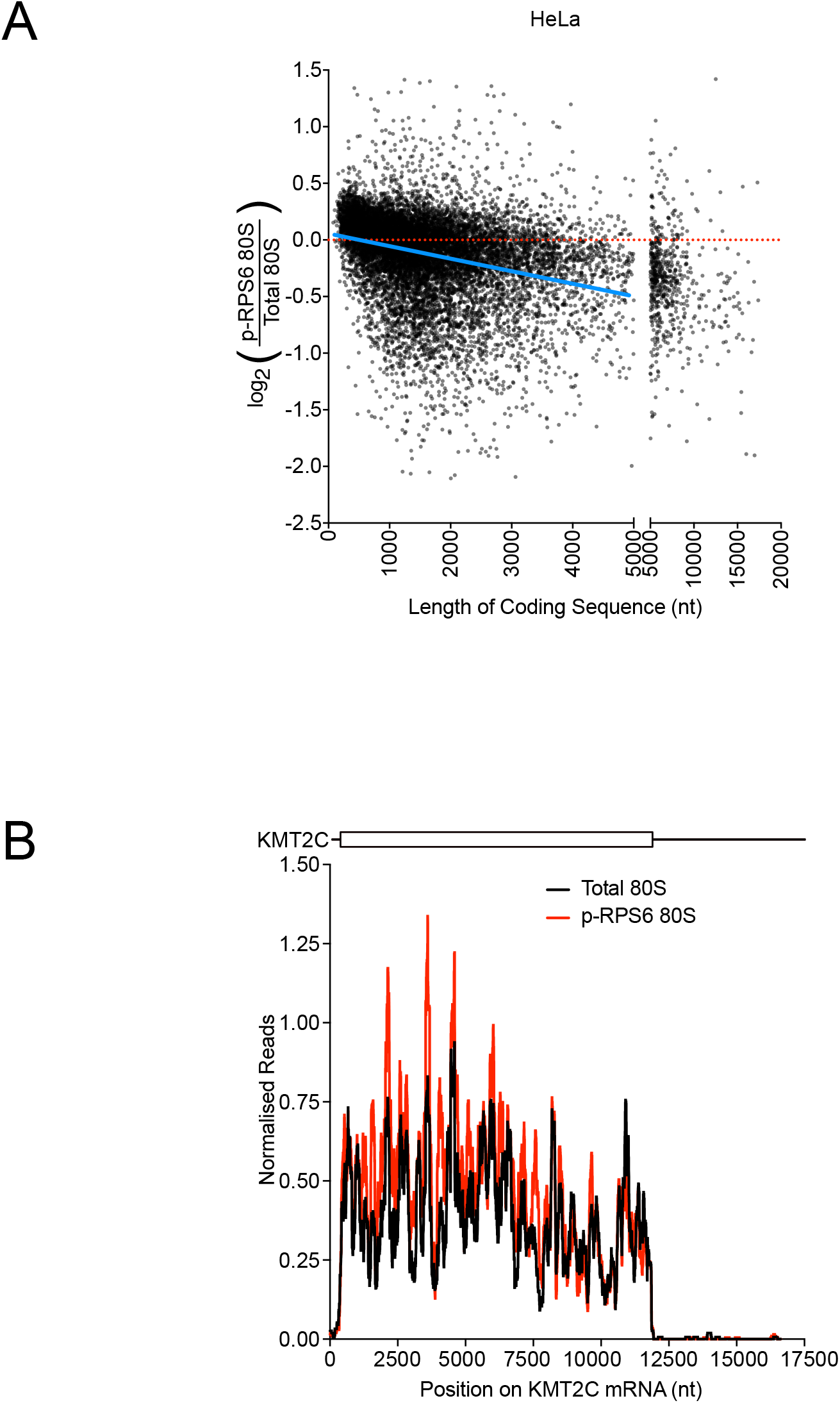
Support to Main Figure 3. **(A)** Scatter plot for relative pS6 abundance (pS6 reads normalized to total 80S reads) versus CDS length for all detected transcripts in HeLa cells. **(B)** Ribosomes translating the mRNA encoding KMT2C (a nuclear protein) are only weakly dephosphorylated on RPS6 Serine 235 and 236. Ribosome occupancy of total 80S and p-RPS6 80S ribosomes along the KMT2C mRNA in HeLa cells. Curves are normalized to sequencing depth. Sliding window smoothening of 150 nt was applied to the curves.

**Suppl. Figure 3:**
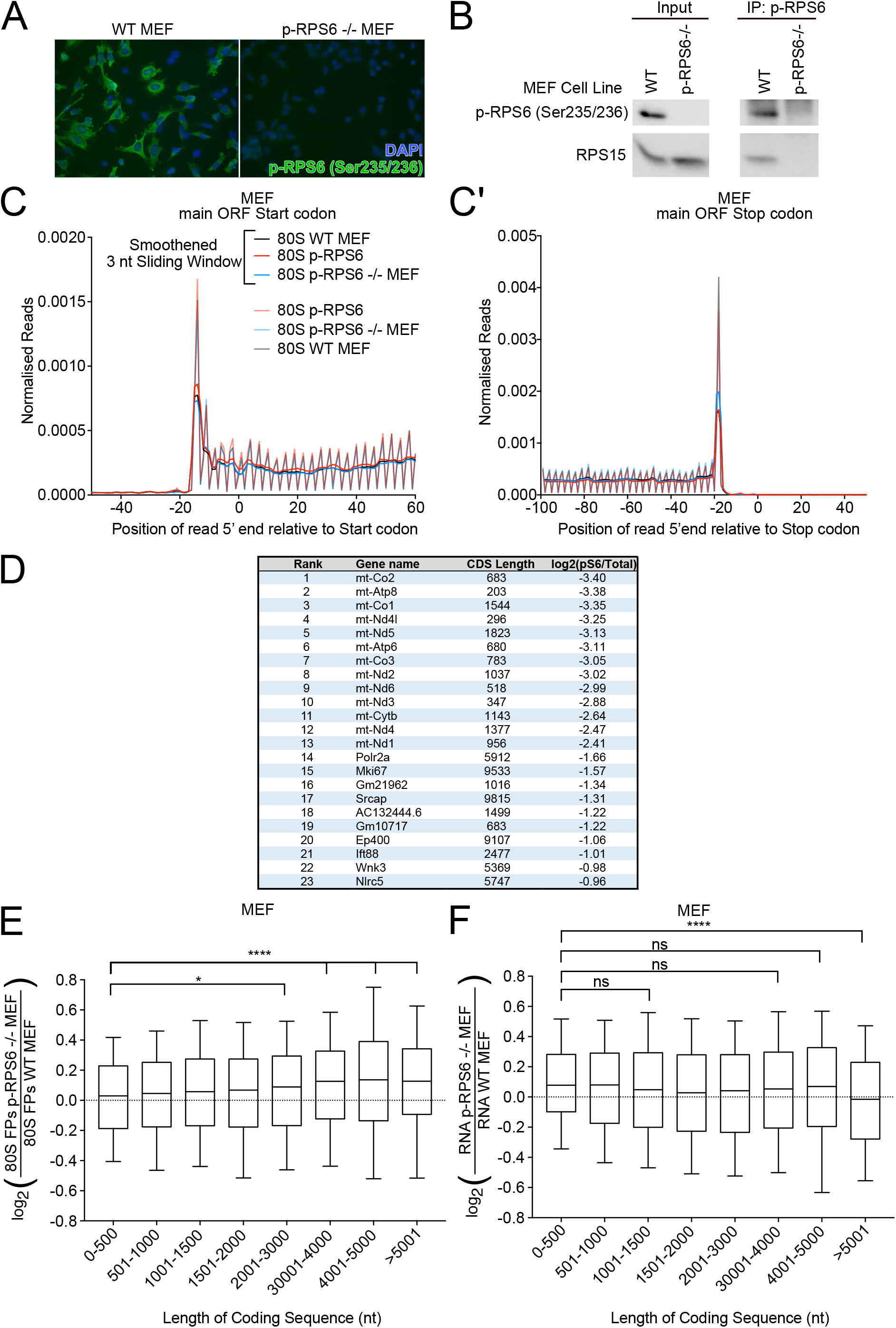
Characterization of rpS6^P−/−^ MEFs and quality controls for total and selective ribosome footprinting. **(A)** rpS6^P−/−^ MEFs lack RPS6 phosphorylation, detected by immunofluorescent staining. Cells were stained for p-RPS6 on Serine 235 and Serine 236 (green) and for DNA with DAPI (blue). **(B)** Immunoprecipitation of ribosomes containing phosphorylated RPS6 is specific in mouse embryonic fibroblasts. Upon immunoprecipitation with p-RPS6 antibody, no ribosomal proteins were detected in the IP from rpS6^P−/−^ MEFs. **(C-C’)** Quality control of p-RPS6-selective and total 80S ribosome footprinting in MEF cells. Metagene profiles for total and p-RPS6-selective 80S ribosome footprints from MEF cells showing the position of the 5’end of ribosome footprints relative to main ORF (C) start and (C’) stop codons for all transcripts. Read counts were normalized to sequencing depth. “Smoothened” indicates the curve was smoothened with a 3nt sliding window. **(D)** As expected, mRNAs with the lowest phospho-S6 abundance in MEFs are mitochondrially encoded and translated. **(E-F)** rpS6^P−/−^ MEF cells have fewer ribosome footprints and not higher mRNA levels on mRNAs with short coding sequences. Grouped analysis of the relationship between coding sequence length and (E) ribosome footprint number or (F) RNA levels in rpS6^P−/−^ MEFs relative to control MEFs. Genes were classified into the indicated groups depending on their annotated coding sequence length. The middle line denotes the median, boxes indicate the upper and lower quartile boundaries and whiskers indicate the first and last decile boundaries. P-values were determined using unpaired, two-sided Kruksal Wallis tests, p-values were adjusted for multiple testing using statistical hypothesis testing, *p<0.0332, ****p<0.0001

**Suppl. Figure 4:**
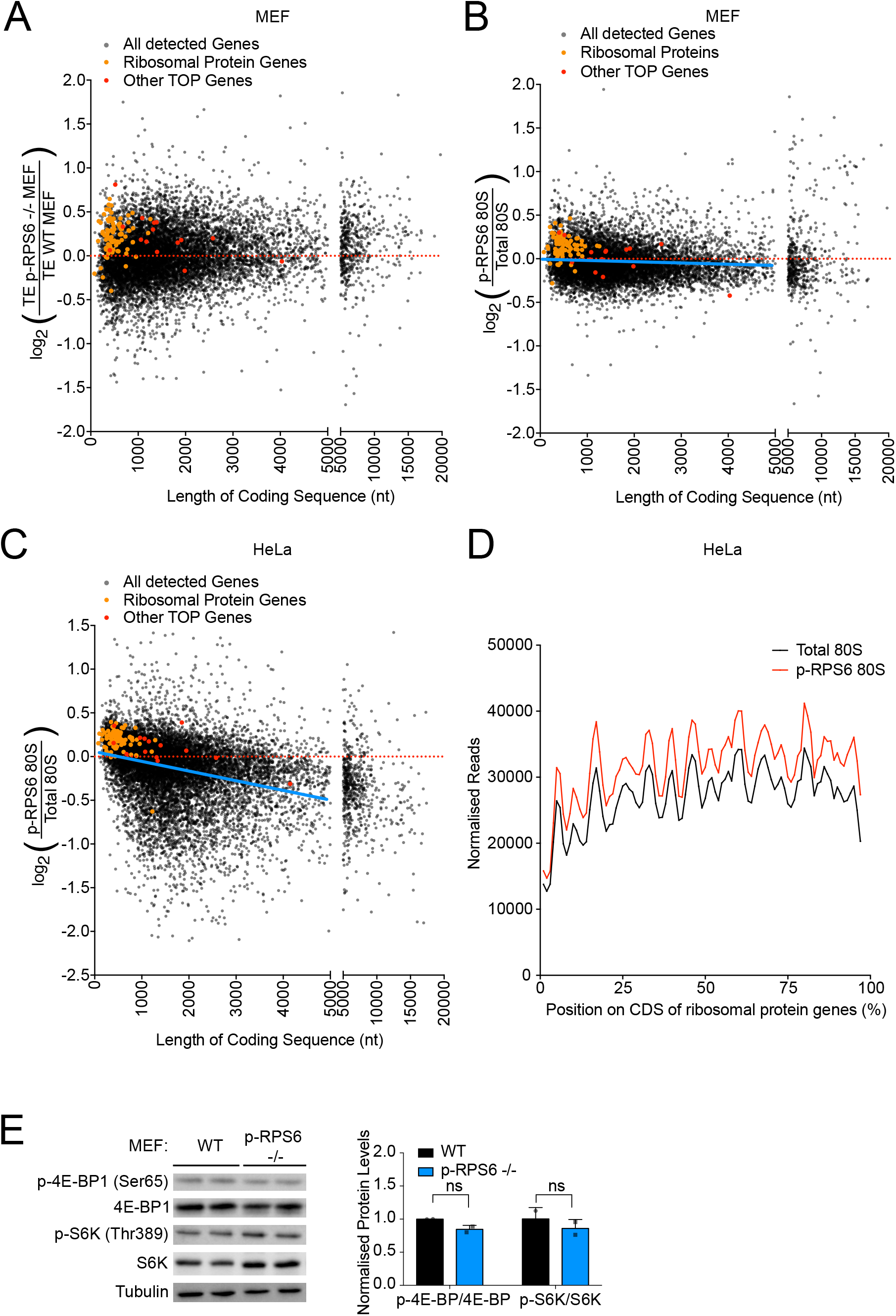
Ribosomal protein and TOP-containing mRNAs escape the translational repression observed for other mRNAs containing short CDSs in rpS6^P−/−^ MEFs. **(A)** Ribosomal protein and TOP-motif containing mRNAs are more efficiently translated in rpS6^P−/−^ MEFs compared to mRNAs of similar length. Scatter plot of log2(*Δ*TE) between rpS6^P−/−^ and control MEF cells versus coding sequence length for all detected genes. Ribosomal protein and TOP genes are specifically indicated. **(B-C)** Ribosomal protein and TOP-motif containing mRNAs are translated by ribosomes containing phosphorylated RPS6 in MEF (B) and HeLa cells (C). Scatter plot of log2(p-RPS6 Enrichment) versus coding sequence length for all detected genes. Ribosomal protein and TOP genes are specifically indicated. Blue lines indicates the correlation between CDS length and average pS6-ribosome occupancy for mRNAs with CDS up to 5000nt long. **(D)** mRNAs encoding ribosomal proteins are translated by ribosomes highly phosphorylated on RPS6 Serine 235 and 236, distributed homogeneously across the coding sequence. Metagene plot of footprints on ribosomal protein mRNAs (n=84). Position along CDS is scaled from 0% (start codon) to 100% (stop codon). Curves normalized to sequencing depth. **(E)** mTOR signaling is not altered in rpS6^P−/−^ MEFs. (left) Immunoblot analysis of two biological replicates of wildtype control and rpS6^P−/−^ MEFs to detect two direct mTORC1 targets, pS6K and p4E-BP. (right) Quantification of p-4E-BP and p-S6K relative total protein levels. Statistical significance of differences was tested with two-sided, unpaired t-tests.

**Suppl. Figure 5:**
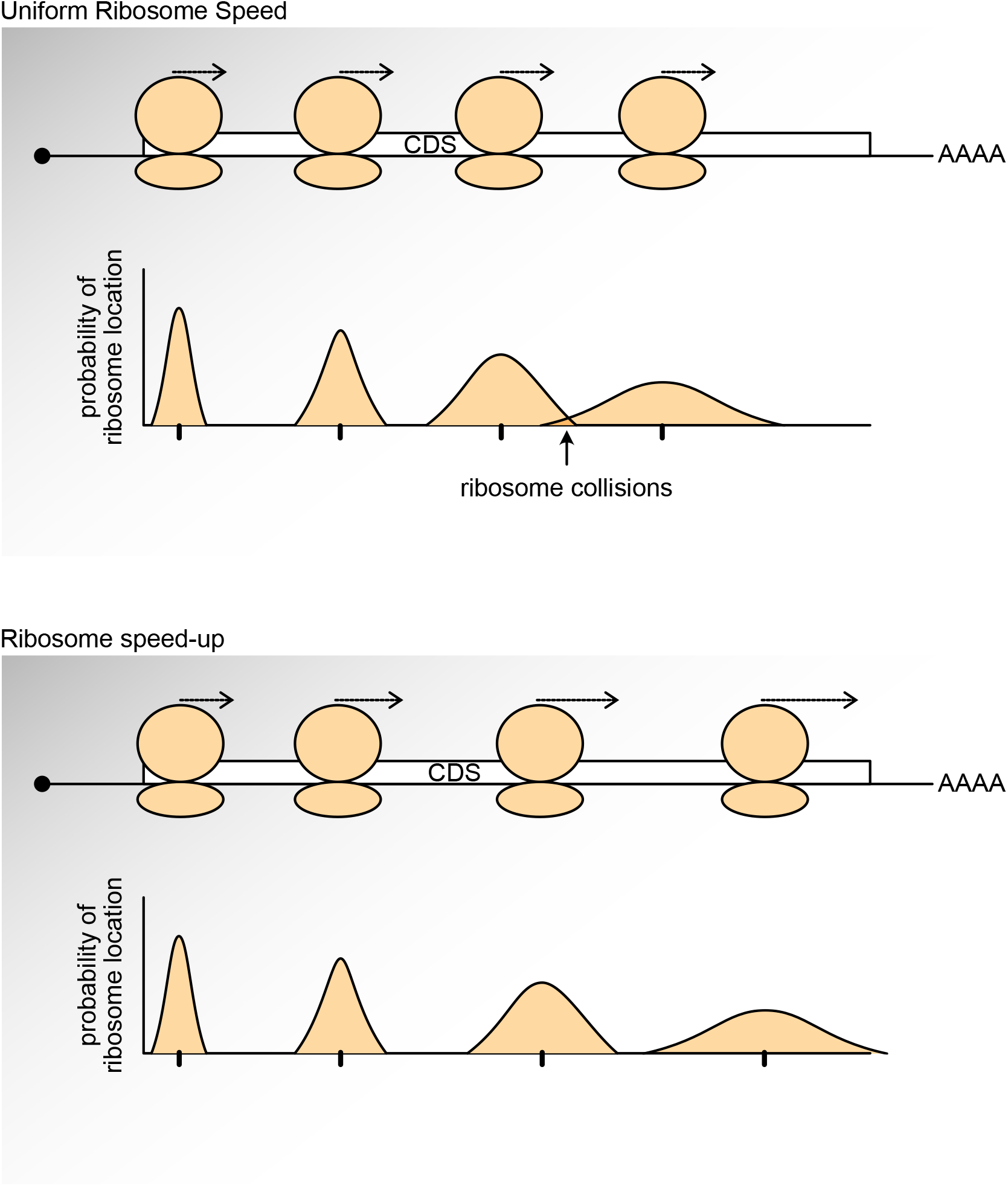
Schematic diagram illustrating how ribosome elongation rate acceleration can prevent ribosome collisions caused by stochastic events. As a ribosome elongates, stochastic events cause it to randomly speed up or slow down compared to the ribosome ‘behind’ it. After many elongation steps this can lead to ‘dephasing’ of the two ribosomes and collisions. Acceleration of ribosome elongation can compensate for this.

## Notes

https://www.ncbi.nlm.nih.gov/geo/query/acc.cgi?acc=GSE168977

### Competing Interest Statement

The authors have declared no competing interest.

